# The effect of context congruency on fMRI repetition suppression for objects

**DOI:** 10.1101/2022.11.17.516972

**Authors:** Chenglin Li, Linda Ficco, Sabrina Trapp, Sophie-Marie Rostalski, Lukas Korn, Gyula Kovács

## Abstract

The recognition of objects is strongly facilitated when they are presented in the context of other objects (Biederman, 1972). Such contexts facilitate perception and induce expectations of context-congruent objects (Trapp & Bar, 2015). The neural mechanisms underlying these facilitatory effects of context on object processing, however, are not yet fully understood. In the present study, we investigate how context-induced expectations affect subsequent object processing. We used functional magnetic resonance imaging and measured repetition suppression, a proxy for prediction error processing, for pairs of alternating or repeated object images, preceded by context-congruent, context-incongruent or neutral cues. We found a stronger repetition suppression in congruent as compared to incongruent or neutral cues in the object sensitive lateral occipital cortex. Interestingly, this effect was driven by enhanced responses to alternating stimulus pairs in the congruent contexts. In addition, in the congruency condition, we discovered significant functional connectivity between object-responsive and frontal cortical regions, as well as between object-responsive regions and the fusiform gyrus. Our findings unravel the neural mechanisms underlying context facilitation.

## Introduction

Objects usually coexist with other objects in our environment, for instance, a hairdryer usually co-occurs with a towel, a toothbrush and other items in a bathroom. Such items compose scenes with specific contextual relations (Kim & Biederman, 2011^1^). Accordingly, previous studies have found that objects are recognized more rapidly and accurately when they are encountered in a congruent compared to incongruent contexts (e.g., Bar & Aminoff, 2003; Biederman, 1972; Chun, 2000; Henderson & Hollingworth, 1999; Kaiser et al., 2019; Oliva & Torralba, 2007). The neural correlates of the perception of context have been studied extensively in the past (e.g., Bar & Aminoff, 2003; Brandman & Peelen, 2017; Caplette et al., 2020; Goh et al., 2004; Hamm et al., 2002; Jenkins et al., 2010; McAndrews et al., 2016; Remy, Saint-Aubert, et al., 2013; Remy, Vayssiere et al., 2014; Remy et al., 2020). Since congruent contexts may induce expectations regarding the occurrence of a given object (Bar, 2004; Trapp & Bar, 2015), the context-based facilitation of object recognition was recently explained by predictive theories. In principle, congruency between a context and an object might reduce the prediction error. Thus, as a consequence, neural responses should be reduced for congruent when compared to incongruent objects (Kok & de Lange, 2015). However, the extent to which context congruence modulates the neural correlates of prediction error has not yet been explicitly investigated.

One way to directly investigate neural correlates of prediction error processing is probing for correlates of neural repetition suppression (RS). Repetition suppression^2^, i.e., the reduction of neural responses for repeated presentations of a given stimulus (Desimone, 1996), has been interpreted as the reduction of prediction error by various researchers (for a review see Grotheer & Kovács, 2016). Although the exact neural mechanisms behind RS remain unclear (Grill-Spector et al., 2006), RS is a very robust neural phenomenon and is widely used to localize the effects of visual category perception, both in the visual and the auditory domain. Importantly, RS seems to be modulated by stimulus expectation. In a seminal study, Summerfield et al (2008) found that the magnitude of RS is modulated by the probability of stimulus repetitions: In the fusiform face area (FFA; Kanwisher et al., 1997), RS was enhanced when faces were presented in blocks in which repetitions were more frequent (and thereby expected) as compared to blocks with repetitions having a low probability (unexpected). The authors suggested that probability-based expectations modulate repetition suppression effects. They interpreted the larger RS in expected blocks as a correlate of reduced prediction error processing (e.g., Henson, 2003; D’Astolfo & Rief, 2017; Kok & de Lange, 2015; Sutton & Barto, 2018).

Thus, on one hand, context-congruent objects lower the neural response as compared to context-incongruent ones. On the other hand, RS is a proxy for prediction error processing, and a reduced RS is the index for lower prediction error processing. If contextual information indeed reduces prediction error processing, one would hypothesize that an object-pair following a congruent context would lead to larger RS, when compared to incongruent or uninformative contexts.

In the past, most neuroimaging studies that investigated context-congruency effects, used objects superimposed on images of scenes (e.g. presenting a rectangular image of a hairdryer in the middle of a larger scene background image; Brandman & Peelen, 2017; Goh et al., 2004; Hamm et al., 2002; Jenkins et al., 2010; McAndrews et al., 2016; Remy, Saint-Aubert, et al., 2013; Remy, Vayssiere et al., 2014; Remy et al., 2020; but see a sequential context-object design in Caplette et al., 2020). Despite its appeal, the simultaneous presentation of context and object images has important limitations. First, one could argue that simply superimposing object images on scenes creates artificial composite images that lack ecological validity. Second, such composite stimuli render it hard to evaluate the context-object interactions or modulations. For instance, the context-congruent effects in object processing could originate from matching processes of context and object, but also from low-level interactions of objects, such as crowding (Caplette et al., 2020). Third, looking at the difference between brain responses in congruent and incongruent conditions, it is difficult to draw conclusions about the underlying mechanism of the context modulations and on prediction error processing directly.

In the current study, we performed a functional magnetic resonance imaging (fMRI) experiment with a novel paradigm that allows us to test context modulation on object processing, while addressing the above-mentioned limitations. To this end, we first presented context images as cues, followed by either repeated or alternated images of objects, which could either be congruent, incongruent or neutral with regard to the prior context cues. This paradigm enables the measurement of RS, while separating the context and object related neural responses. Our findings suggest that the modulation of context on object recognition is mainly driven by the violation of expectations in response to context-incongruent and thereby surprising stimuli. We discuss our results with reference to the theory of predictive coding (PC).

## Materials and Methods

### Participants

Twenty-eight healthy participants took part in the experiment. The sample size, which was determined a priori, ensured at least 80% power and an alpha level of 0.05 to detect within-subject experimental effects with a medium effect size (f > 0.25) (G*Power 3.1.9.7; Faul et al., 2007). Two participants were excluded from the final analysis due to excessive motion during the measurements. The remaining 26 participants (15 females; mean age = 22.7 years; SD = 3.91 years; two left-handed) had normal or corrected-to-normal vision. They received monetary compensation or a 3D-printed model of their own brain. All participants were informed about the experimental procedures and provided their written informed consents before joining the study. No participant had a history of neurological disorders. The experimental protocol was approved by the ethics committee of the Friedrich-Schiller-Universität Jena and conducted in accordance with the guidelines of the Declaration of Helsinki.

### Stimuli

To render the object images from different contexts comparable, we selected objects that were equally plausible for different contexts, and all stemmed from each context similarly. At the same time, they were also subjectively sufficiently distinct. For example, there are exemplars for the category “bag”, which are typical for each of our contexts but at the same time look substantially different in each context (for stimulus examples see Figure 1). Therefore, the final matrix had five contexts (*“kindergarten”, “forest”, “beach”, “hospital”, and “urban life”*) and four stimulus categories (*“chairs”, “bags”, “tools”, and “vehicles”*). For every object category of a given scene nice colored images were downloaded from the public domain of the internet, using the Google image search engine. In addition, to ensure that participants maintained their attention during the fMRI measurement, we added six images of butterflies as target stimuli, which participants were instructed to detect by pressing a button. These target trials were later modelled explicitly but excluded from further analyses. The background of object images was removed by the remove.bg software (remove.bg for Windows 1.4.2). All images were resized to 400 x 400 pixels by GIMP (GNU Image Manipulation Program). At the end of the fMRI experiment, participants performed a sorting task, where they had to assign each object image to one of the five categories. The results showed a high accuracy for allocating the object images into their respective contexts (mean = 90.0%, SE = 2.55%), showing that the objects were diagnostic of their selected contexts.

**Figure 1.**
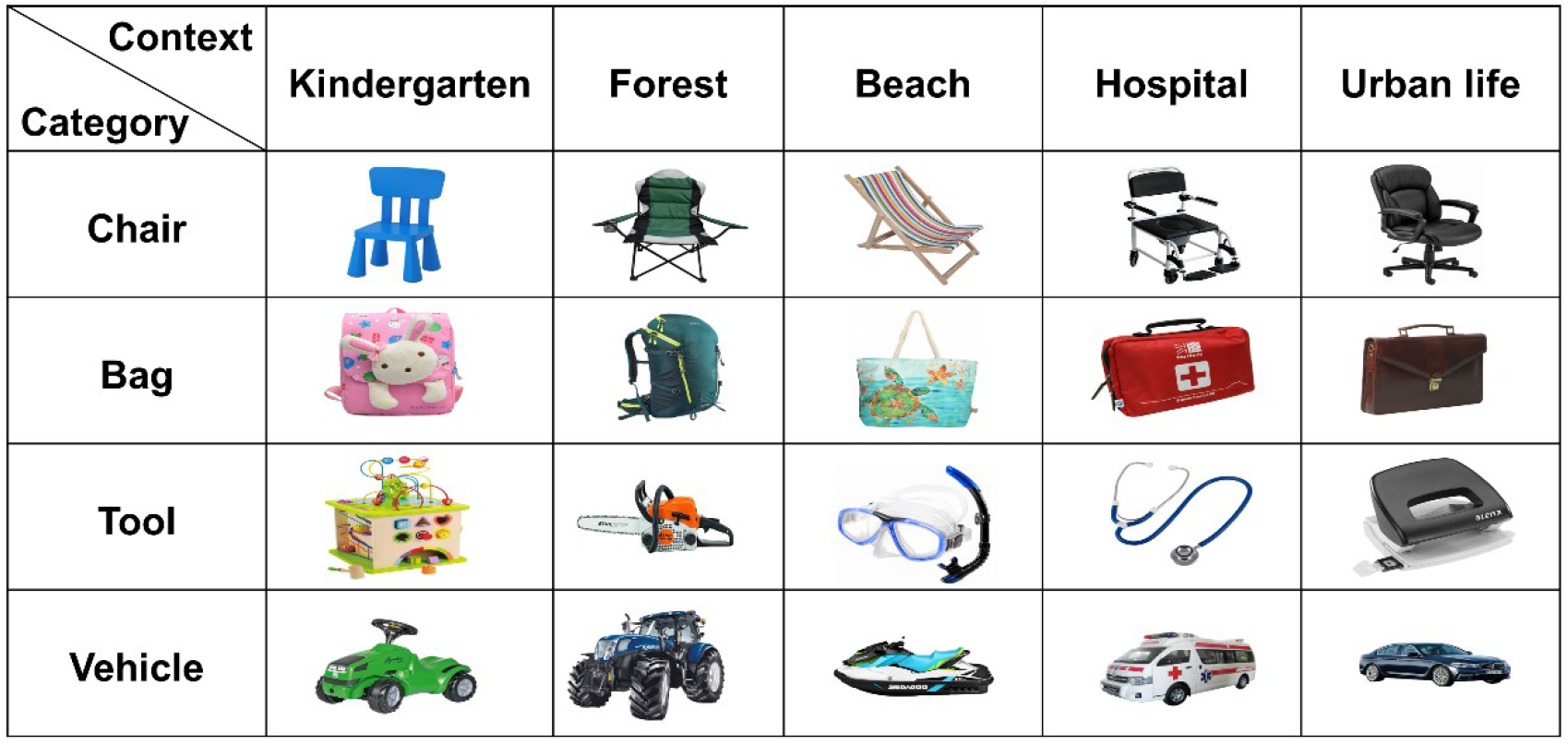
Examples of the object stimuli, separately for the five visual scenes and four categories.

To induce strong contextual effects, two images of a characteristic visual scenery, e.g., a hospital room with beds, were collected for each context to serve as a cue. In addition, two randomized visual noise images were used as neutral cues (Fig 2). Thus, in total, 12 images were used as the contextual cues (size = 1080 x 720 pixels).

**Figure 2.**
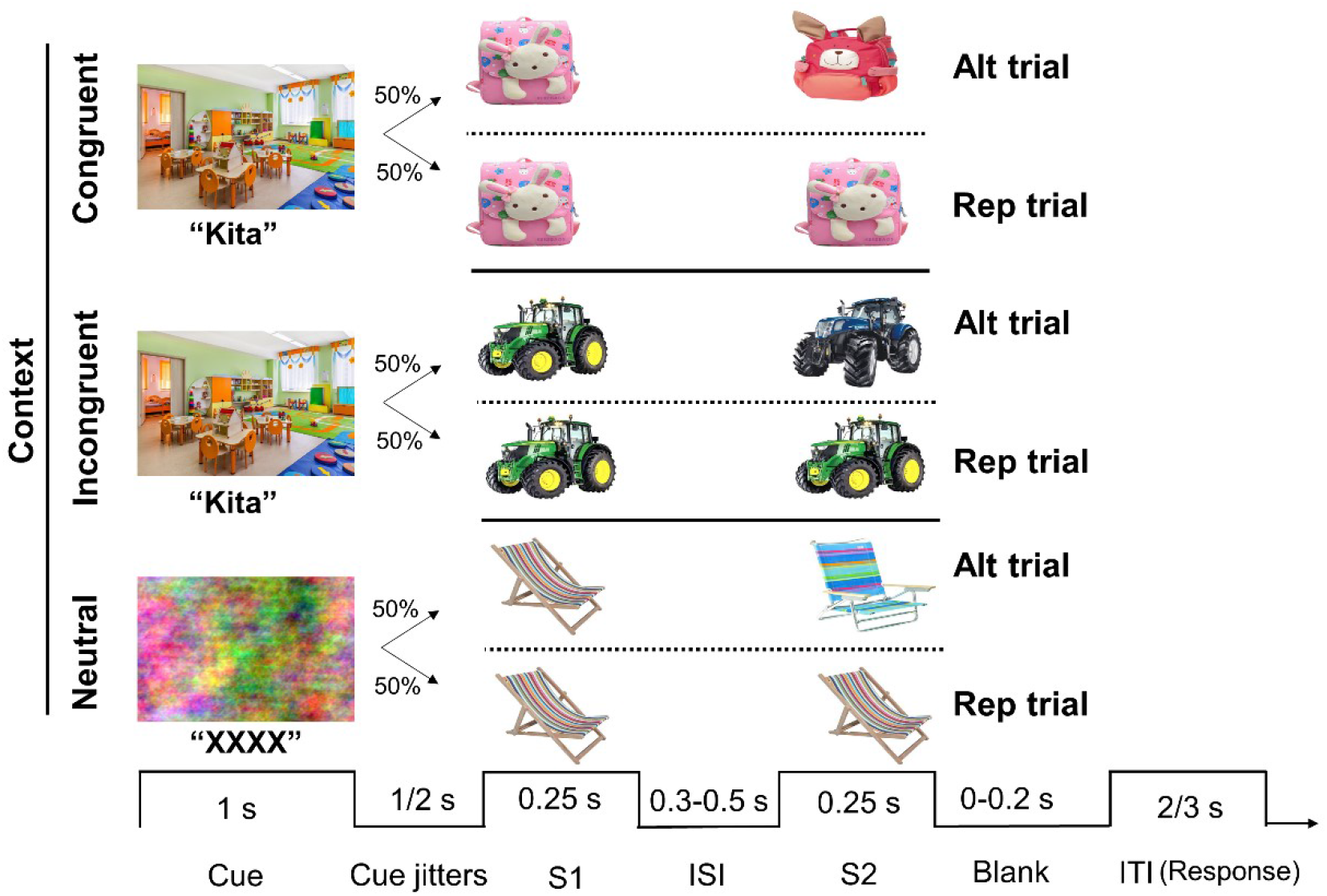
Schematic illustration of the experimental design and trial structure. Altalternation, Rep-repetition, S1-the first stimulus, S2-the second stimuli, ISI-interstimulus-interval, ITI-inter-trial-interval.

### Experimental design and procedure

We adopted a classic stimulus repetition paradigm to measure RS to objects followed by either congruent, incongruent, or neutral cues. The trial structure and the paradigm are illustrated on Figure 2. Psychtoolbox (Version 3.0.15) was used for stimulus presentation and behavioral response collection, conducted under Matlab (R2013a; The MathWorks, Natick, MA, USA).

The fMRI measurement consisted of three runs, each including 132 trials (120 nontarget and 12 target trials). Non-target trials included six different trial types: alternating (Alt) and repeated (Rep) object image-pairs following congruent contexts, Alt and Rep object image-pairs following incongruent contexts and Alt and Rep object image-pairs following the neutral, no-context cues (20 trials per condition, presented randomly). Alt and Rep trials appeared with equal probability, randomly. Each trial commenced with a contextual cue image, presented along its written label (for example “Kita” (kindergarten in German)) or a noise image with the label “XXXX” in the neutral condition. This cue was presented centrally for 1,000 ms and was randomly followed by an inter-stimulus interval of 1,000 or 2,000 ms. Next, a stimulus-pair was presented (exposition time: 250 ms for both), separated by an inter-stimulus-interval (ISI) of 300-500 ms (randomized across trials), and followed randomly by an Inter-trial-interval (ITI) of 2,000 ms or 3,000 ms. The stimulus pairs depicted either the same image (Rep trial) or two different objects belonging to the same context (Alt trials). In target trials, a butterfly image appeared randomly as the first or second stimulus and participants were required to press a button when detecting it. These target trials had the sole purpose of ensuring participants’ attention during task execution and have been excluded from the statistical analyses.

### Imaging parameters and data analysis

The experiment was performed using a 3-Tesla MR scanner (Siemens MAGNETOM Prisma fit, Erlangen, Germany) with a 64-channel head coil. An MP-RAGE sequence was used to acquire a T1-weighted high-resolution 3D anatomical image (192 slices; TR = 2300 ms; TE = 3.03 s; flip angle = 90°; 1 mm isotropic voxel size). During the functional runs, to obtain a temporal resolution of 1 second, T2*-weighted fMRI-images were collected with a multi-band EPI sequence (MB acceleration factor = 8) under the following parameters: 64 slices; FOV = 200 x 2000 mm^2^; TR = 1000 ms; TE = 45.8 ms; flip angle = 90°; 2 mm isotropic voxel size. Data were preprocessed using SPM12 (Welcome Department of Imaging Neuroscience, London, UK). Briefly, the functional images were slice-timed, realigned, and co-registered to structural scans. The functional images were then normalized to the MNI-152 space, resampled to 1 x 1 x 1 mm resolution, and spatially smoothed using a 6-mm Gaussian kernel.

An additional functional localizer run was conducted to define regions of interest (ROIs). Scenes, objects, and Fourier-randomized noise images were presented (4 Hz; 230ms exposition time; 20 ms ISI) in blocks of 10 s, interrupted by breaks of 10 s and repeated five times. Each block included 40 images (size: 400 x 400 pixels with a grey background). Participants performed a passive viewing task during these localizer runs. Like prior studies (Brandman & Peelen, 2017; Jenkins et al., 2010; Remy et al., 2020), we focused on the lateral occipital complex (LOC; Malach et al., 1995), a key region of the ventral visual pathway relevant for object processing (for a review see Grill-Spector et al., 2001). The location of LOC was determined individually by contrasting object and noise blocks and established as the local maximum from the t-maps with a threshold of *P*_FWE_ < 0.05 on the single-subject level. The average MNI coordinates (± SE) were 42.5 (1.07), −78.0 (1.16), −3.2 (1.66) for the right LOC and −42.3 (0.95), −81.0 (1.09), −3.7 (1.42) for the left LOC, respectively. Canonical hemodynamic response functions were extracted using MarsBaR 0.44 toolbox for SPM 12 (Brett et al., 2002). The peak BOLD values were extracted from the event-related runs and analyzed by repeated-measures ANOVAs with hemisphere (2), contextual congruence (3, congruent, incongruent, and neutral) and trial type (2, Alt and Rep) as within-subject factors. All multiple comparisons of post-hoc tests were corrected with the Fisher’s method.

Task-based functional connectivity analyses were performed using the CONN toolbox (version 21.a; Whitfield-Gabrieli et al., 2012) as implemented in Matlab 2018a. After importing the preprocessed fMRI data, we denoised it using the component-based noise correction (CompCor) method to remove noise components arising from white matter and cerebrospinal areas, as well as participants’ motion parameters, and bandpass filtered it (0.01 - Inf). We performed a generalized Psychophysiological Interactions (gPPI) analysis implemented in CONN toolbox (McLaren et al., 2012). In the first-level connectivity analyses, the right and left LOC were used as the seed for seed-to-voxel functional connectivity. At the second-level, we adopted custom contrasts to compute the main effects of contextual congruence, trial type and their interaction, respectively. Results were thresholded at *p* < 0.05 (voxelwise, uncorrected), with a cluster correction for multiple comparisons at FDR < 0.05.

## Results

### Behavioural results

Participants detected the target stimuli on average with 96.0% (±SE: 1.83%) accuracy during the fMRI session, suggesting that they paid attention to the stimuli during the sessions.

### Repetition suppression in the LOC

We found a significant main effect of trial type (*F*(1, 25) = 107.318, *p* < 0.001, *η_p_^2^* = 0.811), with lower BOLD signals in the Rep (0.498 ± 0.056) when compared to Alt trials (0.615 ± 0.062) (Figure 3). In addition, there were significant main effects of congruence (*F*(2, 50) = 13.754, *p* < 0.001, *η_p_^2^* = 0.355) and hemisphere (*F*(1, 25) = 7.077, *p* = 0.013, *η_p_^2^* = 0.221), with larger responses in the congruent condition and generally a higher activity level over the right hemisphere. Importantly, we found a significant interaction between trial type and congruence (*F*(2, 50) = 4.490, *p* = 0.016, *η_p_^2^* = 0.152) suggesting a modulation of RS by contextual congruency. Post-hoc tests revealed stronger RS in the congruent (Fisher LSD *post hoc* test: *p* = 0.0000000314) compared to the incongruent (Fisher LSD *post hoc* test: *p* = 0.000171) or neutral (Fisher LSD *post hoc* test: *p* = 0.024) condition. Interestingly, the higher RS magnitude was driven by the larger responses in the Alt trials in both congruent and incongruent conditions, as suggested by the post-hoc tests of the Alt trials for the Congruent vs Incongruent (Fisher LSD *post hoc* test: *p* < 0.011) and Congruent vs Neutral conditions (Fisher LSD *post hoc* test: *p* = 0.000000025), while the Rep trials were similar across all three context conditions (Fisher LSD *post hoc* test: *p* > 0.05 for each condition). No other interaction effects reached significance.

**Figure 3.**
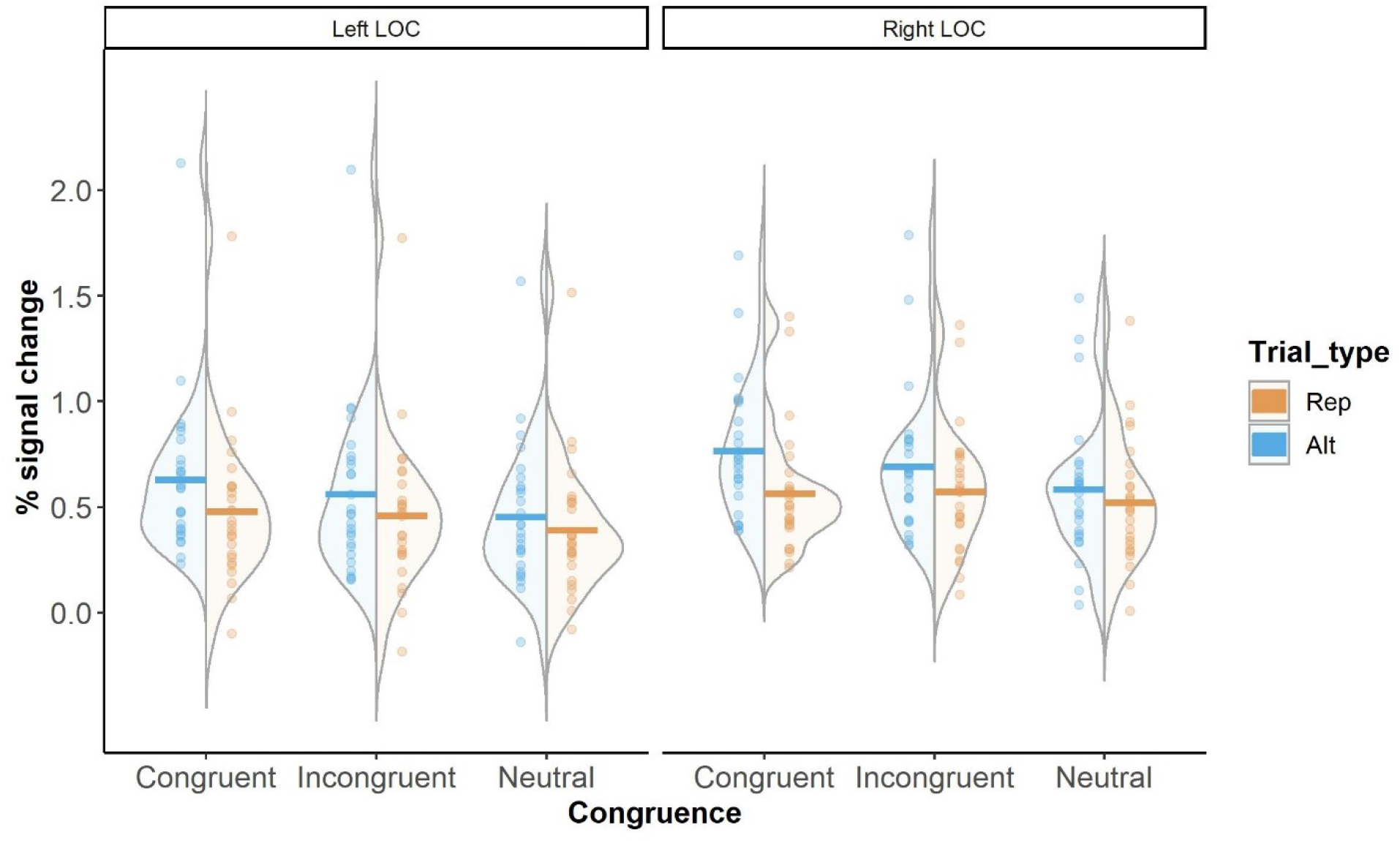
The modulation of context on RS for objects in the LOC. Activation profile of the left and right LOC for Alt and Rep trials in the congruent, incongruent and neutral conditions, separately. Horizontal crossbars represent the mean signal, whereas points represent individual participant’s data.

### Task-based functional connectivity during repetition suppression

To study the origins of the RS effect and its context modulation, we explored functional connectivity. The left and right LOCs served as seeds. We investigated the main effect of trial type (RS), and the main effect of congruence (Congruence), as well as their interaction effect separately. The results are presented in Figure 4 and Table I. For RS, we found strong connectivity between the right LOC and the right frontal pole and middle frontal gyrus (*t*(25) > 2.06, *p* ≥ 0.015_FDR-corrected_). However, when we used the left LOC as seed, significant connectivity was found with the right posterior supramarginal gyrus (*t*(25) > 2.49, *p* > 0.005_FDR-corrected_). For the congruence effect, similarly we found significant connectivity between the right LOC and the left frontal pole (*F*(2,50) > 5.90, *p* > 0.007_FDR-corrected_), and between the left LOC and the left posterior supramarginal gyrus (*F*(2,50) > 5.90, *p* > 0.012_FDR-corrected_). Finally, for the trial x context interaction effect, significant connectivity was found between the right LOC and the fusiform gyrus bilaterally, (*F*(2,50) > 3.18, *p* ≥ 0.025_FDR-corrected_).

**Figure 4.**
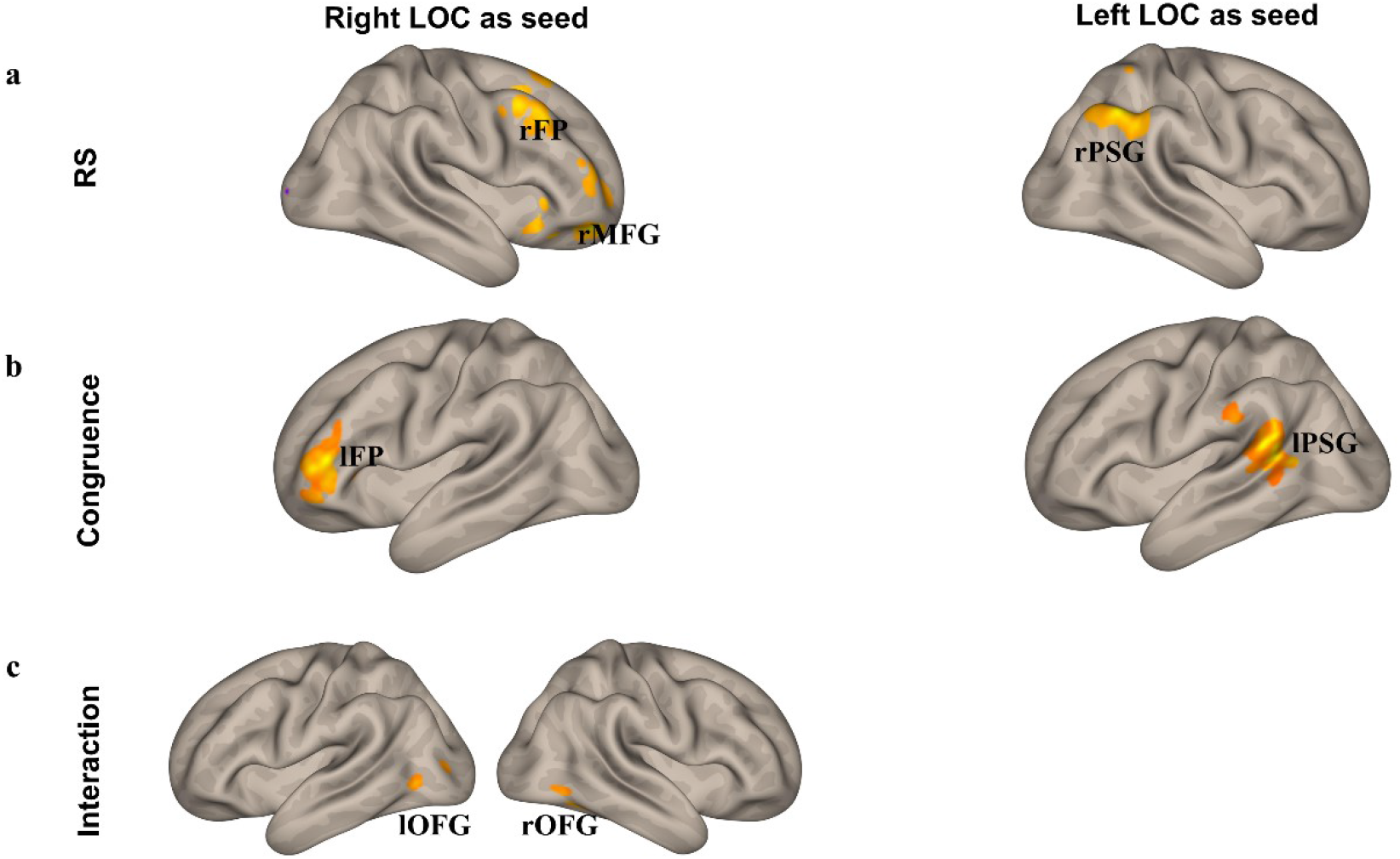
Functional connectivity results with the right and left LOC as seed for the main effect of trial type (RS; a), and the main effect of congruence (Congruence; b), as well as their interaction effect (Interaction; c) separately. rFP – right frontal pole, rMFG – right middle frontal gyrus, rPSG – right posterior supramarginal gyrus, lFP – left frontal pole, lPSG – left posterior supramarginal gyrus, lOFG – left occipital fusiform gyrus, rOFG – right occipital fusiform gyrus.

**Table 1.** Functional connectivity results with the LOC seed. rLOC – right lateral occipital complex, lLOC – left lateral occipital complex.

| Effect | Seed | Cluster<br>(MNI coordinates) | Cluster<br>size<br>(voxels) | Region | *P* (FDR-corrected) |
| --- | --- | --- | --- | --- | --- |
| RS | rLOC | 30, 52, -14 | 586 | Right frontal pole | 0.015 |
|  | rLOC | 42, 12, 44 | 556 | Right middle frontal gyrus | 0.016 |
|  | lLOC | 48, -50, 54 | 838 | Right posterior supramarginal gyrus | 0.005 |
| Congruence | rLOC | -32, 38, 4 | 158 | Left frontal pole | 0.007 |
|  | lLOC | -64, -56, 20 | 159 | Left posterior supramarginal gyrus | 0.012 |
| Interaction | rLOC | 36, -48, -8 | 764 | Right occipital fusiform gyrus | 0.005 |
|  | rLOC | -32, -74, 4 | 550 | Left occipital fusiform gyrus | 0.025 |
|  | lLOC | - | - | - | - |

## Discussion

The major results of the current study are the following: (1) We found stronger RS for congruent than for incongruent or no context conditions; (2) the LOC shows functional connectivity with frontal and parietal regions, and these connections are influenced by stimulus repetition and context-object congruency; (3) the LOC appears to be functionally connected to the fusiform gyrus as well, specifically for the modulation of RS by stimulus context.

In the current study, we adopted a new paradigm to investigate the modulation of RS as a proxy for prediction error processing by context. Previous neuroimaging studies addressed contextual processing by comparing brain activity either when participants viewed isolated objects, or with objects superimposed artificially on congruent and incongruent context images (e.g., Brandman & Peelen, 2017; Goh et al., 2004; Hamm et al., 2002; Jenkins et al., 2010; McAndrews et al., 2016; Remy, Saint-Aubert, et al., 2013; Remy, Vayssiere et al., 2014; Remy et al., 2020). In addition, some studies adopted a block design, where participants were presented with objects in isolation, but in congruent or incongruent context blocks (e.g., Bar & Aminoff, 2003) or pairs (e.g., Livne & Bar, 2016). Some of these studies found significant and wide-spread activation differences for context incongruent compared to context congruent objects (Faivre et al., 2019; Remy et al., 2014; 2020). However, other studies reported the exact opposite, i.e, (Crafa et al., 2017; McAndrews et al., 2016; van Kesteren et al., 2020) and others even showed no difference at all (Caplette et al., 2020). As compared to previous studies, one of our aims was to improve the ecological validity of the stimulation. In fact, contexts are typically processed *prior* to objects (Bar, 2003; Oliva & Torralba, 2007; Caplette et al., 2020). At the same time, our approach is better suited for disentangling context-object interactions and context-based predictions, as any effects found in simultaneous paradigms could be due to lower level perceptual matching processes or crowding effects (Caplette et al., 2020). In addition, we included a neutral condition (for a review see Feuerriegel et al., 2021) in which participants could not form any specific expectations regarding the subsequent objects. Finally, the major aim of our study was to directly look at effects of context information on prediction error processing, using RS as a proxy for reduced or enhanced prediction errors.

Our findings confirm that the strongest RS effect emerges after a congruent, rather than an incongruent or no context, indicating reduced prediction error processing in this condition. However, a surprising aspect of our data is that the context modulation of RS is not driven by a stronger response suppression (i.e. expectation suppression, for a review see Feuerriegel et al, 2021) or a reduced signal to repeated objects (when these are congruent with the context), but by the opposite, i.e., an *enhanced* response to alternating trials during context congruent conditions. In other words, a trial composed of two alternating object images elicits larger responses when it is preceded by a congruent than after an incongruent context cue. Possibly, this could be due to the differential processing of the first and second stimulus of the pair. Thus, after observing a congruency between the context and the *first* object of the stimulus pair, participants may have developed the expectation to see that object again, i.e., when the second stimulus is presented. In the alternating conditions, however, this expectation is violated. As a consequence, the LOC generates a prediction error signal, manifested in the larger BOLD response for the Alt trials. Note that this prediction error signal may be larger in the congruent than in the incongruent condition because here, participants’ contextual expectations are already violated by seeing the first object, thus the presentation of the second object of the pair does not lead to further response enhancements. This hypothesis could also explain the fact that larger RS was observed in the congruent as compared to the neutral condition, because in the later one no contextual expectation is formed at all. This account is supported by previous studies of the predictive modulation of RS that emphasize the contribution of surprise-related response enhancement (e,g., Amado et al., 2016; Kaposvari et al., 2018; for a review see Feuerriegel et al., 2021). For example, Amado et al. (2016) presented pairs of faces that were repeated or alternated. Orthogonally to this, the probability of repetition was manipulated by the gender of the first face (female, male or gender-neutral baby faces) within each pair, thus, repetition or alternation trials could be expected, neutral, or surprising. The BOLD signals in the right FFA showed a larger response for surprising compared to neutral face pairs in the alternation trials, suggesting the critical role of surprise-related response enhancement in the expectation effects.

In recent years, RS has been explained by predictive processing, as repeating a stimulus may induce a reduction of prediction errors (for a review see Grotheer & Kovács, 2016; but see Grill-Spector et al., 2006 for alternative accounts of RS). That, we observed significant RS effects for objects in the LOC, which replicates previous studies (e.g., Grill-Spector et al., 1999; Kourtzi & Kanwisher, 2001; Sawamura et al., 2006). Our functional connectivity analysis shows strong connections between the right LOC and frontal areas of the same hemisphere for the main effect of RS, which supports its top-down accounts, as frontal areas are generally regarded as the potential source of predictions (e.g., Gotts et al., 2015; Summerfield et al., 2006; Ficco et al., 2021). We also found significant functional connectivity between the right LOC and the frontal pole for the context-object congruency effect. This further strengthens the idea that higher-order regions may contribute to top-down expectations which are responsible for the modulation of object processing by contexts (e.g., Bar, 2004; Caplette et al., 2020; Spaak et al., 2022).

Interestingly, most previous studies presenting context and objects simultaneously mainly found interaction effects in the Parahippocampal (PHC) and Retrosplenial (RSC) cortices, areas well-known for their role in scene processing (Bar, 2003, 2004; Bar & Aminoff, 2003; Bar et al., 2006; Livbe & Bar,2016; Brandman & Peelen, 2017). However, our work and that of Caplette et al., 2020, both using sequential context-object presentation paradigms, found context-object interaction effects in frontal and object-sensitive temporal regions. Many factors may account for this difference, the most important being the use of different paradigms. The PHC and the RSC are scene-selective areas (e.g., Brandman & Peelen, 2017; Epstein et al., 1999; Epstein & Kanwisher, 1998; Lescroart & Gallant, 2019), while frontal cortices are related to the generation and violation of predictions (e.g., Gotts et al., 2015; Summerfield et al., 2006; Weilnhammer et al., 2017; for a meta-analysis see Ficco et al., 2021). In simultaneous scene-object presentation paradigms participants view the superimposed image of a context and an object, thereby the context-object interactions might capture context-object congruency or matching processes. Unlike these, sequential paradigms generate expectations in the participants when they see the contextual cues. This might explain why the frontal areas showed a different activation pattern when the context was congruent or incongruent. Again, one can interpret this effect according to the theory of predictive processing (Friston, 2005; Rao & Ballard, 1999): The modulation of context on object processing likely occurs in object-responsive regions after receiving top-down predictions from frontal areas.

One important aspect of the current results is that, when context and repetition interact, the right LOC seems to be functionally connected with the bilateral occipital fusiform gyri. This region, also named as posterior fusiform gyrus, is distinct from the LOC and the rest of the FG (Caspers et al., 2013), is object-selective (Weiner & Zilles, 2016; Weiner et al., 2018) and shows task-based functional connectivity with the LOC (Caspers et al., 2014; Malikovic et al., 2016). A visual comparison between our results (Fig. 4, panel c) and the map reported in Caspers et al., 2014 (Fig. 1, p. 2755) points to the involvement of two different regions of the FG (FG1 and FG2) in our results. These two sub-regions have distinct functional profiles: FG1 shows features of lower-level, whereas FG2 of higher-order regions (Caspers et al., 2014). Both might underpin the interaction between context congruence and stimulus repetition in our study. We speculate that the FG1-like cluster may contribute information on whether low-level features of objects change (as an “alternation detector”), and feed it to the right LOC. Instead, the FG2 in Caspers et al. (2014) seems to be especially involved in reading and processing faces. So, hypothetically, the FG2-like cluster of the current study might process category-diagnostic features of the objects, which would be then integrated in the LOC with context-induced expectations produced in the frontal lobe. However, one has to mention that neither the current connectivity analyses, nor those of Caspers et al. (2014), allow to draw conclusions on the directionality of these connectivity patterns, which can be a promising goal for future studies.

At present, it is unclear why the connectivity profiles of the left and right LOC are different despite their similar activation patterns in response to our stimulus manipulations (cf. Fig. 3). In both conditions, the right LOC is mostly connected to frontal regions, whereas the left LOC connects preferentially to temporo-parietal structures. This is even more surprising, based on evidence of differential activation and connectivity in these two areas depending on task demands (Large et al., 2007; Malikovic et al., 2016). Our findings indicate that the very same functions, i.e. processing object repetition and context congruence, may be accomplished via different networks in the two hemispheres.

In conclusion, the present study shows that the modulation of context on object processing can be observed in the LOC. It also provides evidence about how prediction errors change during context-driven modulation by highlighting the contribution of frontal regions in this process. These findings support the idea that object processing is modulated by top-down contextual expectations.

## Footnotes

1 We dedicate this paper to the memory of Irving Biederman, without whom vision science would not be the same today.

2 Please note that this repetition suppression (RS) effect has also been called stimulus-specific adaptation (Sobotka & Ringo, 1994), or simply as adaptation (Grill-Spector & Malach, 2001).

## References

Amado, C., Hermann, P., Kovács, P., Grotheer, M., Vidnyánszky, Z., & Kovács, G. (2016). The contribution of surprise to the prediction based modulation of fMRI responses. Neuropsychologia, 84, 105–112. https://doi.org/10.1016/j.neuropsychologia.2016.02.003

Bar, M. (2003). A cortical mechanism for triggering top-down facilitation in visual object recognition. Journal of cognitive neuroscience, 15(4), 600–609. https://doi.org/10.1162/089892903321662976

Bar, M., & Aminoff, E. (2003). Cortical analysis of visual context. Neuron, 38(2), 347–358. https://doi.org/10.1016/S0896-6273(03)00167-3

Bar, M. (2004). Visual objects in context. Nature Reviews Neuroscience, 5(8), 617–629. https://doi.org/10.1038/nrn1476

Biederman, I. (1972). Perceiving real-world scenes. Science, 177(4043), 77–80. https://doi.org/10.1126/science.177.4043.77

Brandman, T., & Peelen, M. V. (2017). Interaction between scene and object processing revealed by human fMRI and MEG decoding. Journal of Neuroscience, 37(32), 7700–7710. https://doi.org/10.1523/JNEUROSCI.0582-17.2017

Brett, M., Johnsrude, I. S., & Owen, A. M. (2002). The problem of functional localization in the human brain. Nature reviews neuroscience, 3(3), 243–249. https://doi.org/10.1038/nrn756

Caplette, L., Gosselin, F., Mermillod, M., & Wicker, B. (2020). Real-world expectations and their affective value modulate object processing. NeuroImage, 213, 116736. https://doi.org/10.1016/j.neuroimage.2020.116736

Caspers, J., Zilles, K., Eickhoff, S. B., Schleicher, A., Mohlberg, H., & Amunts, K. (2013). Cytoarchitectonical analysis and probabilistic mapping of two extrastriate areas of the human posterior fusiform gyrus. Brain Structure and Function, 218(2), 511–526. https://doi.org/10.1007/s00429-012-0411-8

Caspers, J., Zilles, K., Amunts, K., Laird, A. R., Fox, P. T., & Eickhoff, S. B. (2014). Functional characterization and differential coactivation patterns of two cytoarchitectonic visual areas on the human posterior fusiform gyrus. Human brain mapping, 35(6), 2754–2767. https://doi.org/10.1002/hbm.22364

Chun, M. M. (2000). Contextual cueing of visual attention. Trends in cognitive sciences, 4(5), 170–178. https://doi.org/10.1016/S1364-6613(00)01476-5

Crafa, D., Hawco, C., & Brodeur, M. B. (2017). Heightened responses of the parahippocampal and retrosplenial cortices during contextualized recognition of congruent objects. Frontiers in behavioral neuroscience, 11, 232. https://doi.org/10.3389/fnbeh.2017.00232

D’Astolfo, L., & Rief, W. (2017). Learning about expectation violation from prediction error paradigms–A meta-analysis on brain processes following a prediction error. Frontiers in psychology, 8, 1253. https://doi.org/10.3389/fpsyg.2017.01253

Desimone, R. (1996). Neural mechanisms for visual memory and their role in attention. Proceedings of the National Academy of Sciences, 93(24), 13494–13499. https://doi.org/10.1073/pnas.93.24.1349

Epstein, R., & Kanwisher, N. (1998). A cortical representation of the local visual environment. Nature, 392(6676), 598–601. https://doi.org/10.1038/33402

Epstein, R., Harris, A., Stanley, D., & Kanwisher, N. (1999). The parahippocampal place area: recognition, navigation, or encoding?. Neuron, 23(1), 115–125. https://doi.org/10.1016/S0896-6273(00)80758-8

Faivre, N., Dubois, J., Schwartz, N., & Mudrik, L. (2019). Imaging object-scene relations processing in visible and invisible natural scenes. Scientific reports, 9(1), 1–13. https://doi.org/10.1038/s41598-019-38654-z

Faul, F., Erdfelder, E., Lang, A. G., & Buchner, A. (2007). G* Power 3: A flexible statistical power analysis program for the social, behavioral, and biomedical sciences. Behavior research methods, 39(2), 175–191. https://doi.org/10.3758/BF03193146

Feuerriegel, D., Vogels, R., & Kovács, G. (2021). Evaluating the evidence for expectation suppression in the visual system. Neuroscience & Biobehavioral Reviews, 126, 368–381. https://doi.org/10.1016/j.neubiorev.2021.04.002

Ficco, L., Mancuso, L., Manuello, J., Teneggi, A., Liloia, D., Duca, S.,… & Cauda, F. (2021). Disentangling predictive processing in the brain: a meta-analytic study in favour of a predictive network. Scientific Reports, 11 (1), 1–14. https://doi.org/10.1038/s41598-021-95603-5

Friston, K. (2005). A theory of cortical responses. Philosophical Transactions of the Royal Society B: Biological Sciences, 360(1456), 815–836. https://doi.org/10.1098/rstb.2005.1622

Goh, J. O., Siong, S. C., Park, D., Gutchess, A., Hebrank, A., & Chee, M. W. (2004). Cortical areas involved in object, background, and object-background processing revealed with functional magnetic resonance adaptation. Journal of Neuroscience, 24(45), 10223–10228. https://doi.org/10.1523/JNEUROSCI.3373-04.2004

Gotts, S. J., Milleville, S. C., & Martin, A. (2015). Object identification leads to a conceptual broadening of object representations in lateral prefrontal cortex. Neuropsychologia, 76, 62–78. https://doi.org/10.1016/j.neuropsychologia.2014.10.041

Grill-Spector, K., Kushnir, T., Edelman, S., Avidan, G., Itzchak, Y., & Malach, R. (1999). Differential processing of objects under various viewing conditions in the human lateral occipital complex. Neuron, 24(1), 187–203. https://doi.org/10.1016/S0896-6273(00)80832-6

Grill-Spector, K., & Malach, R. (2001). fMR-adaptation: a tool for studying the functional properties of human cortical neurons. Acta psychologica, 107(1-3), 293–321. https://doi.org/10.1016/S0001-6918(01)00019-1

Grill-Spector, K., Kourtzi, Z., & Kanwisher, N. (2001). The lateral occipital complex and its role in object recognition. Vision research, 41(10-11), 1409–1422. https://doi.org/10.1016/S0042-6989(01)00073-6

Grill-Spector, K., Henson, R., & Martin, A. (2006). Repetition and the brain: neural models of stimulus-specific effects. Trends in cognitive sciences, 10(1), 14–23. https://doi.org/10.1016/j.tics.2005.11.006

Grotheer, M., & Kovács, G. (2016). Can predictive coding explain repetition suppression?. Cortex, 80, 113–124. https://doi.org/10.1016/j.cortex.2015.11.027

Hamm, J. P., Johnson, B. W., & Kirk, I. J. (2002). Comparison of the N300 and N400 ERPs to picture stimuli in congruent and incongruent contexts. Clinical Neurophysiology, 113(8), 1339–1350. https://doi.org/10.1016/S1388-2457(02)00161-X

Henderson, J. M., & Hollingworth, A. (1999). High-level scene perception. Annual Review of Psychology, 50(1), 243–271. https://doi.org/10.1146/annurev.psych.50.1.243

Henson, R. N. (2003). Neuroimaging studies of priming. Progress in neurobiology, 70(1), 53–81. https://doi.org/10.1016/S0301-0082(03)00086-8

Jenkins, L. J., Yang, Y. J., Goh, J., Hong, Y. Y., & Park, D. C. (2010). Cultural differences in the lateral occipital complex while viewing incongruent scenes. Social cognitive and affective neuroscience, 5(2-3), 236–241.

Kaiser, D., Quek, G. L., Cichy, R. M., & Peelen, M. V. (2019). Object vision in a structured world. Trends in Cognitive Sciences, 23(8), 672–685. https://doi.org/10.1016/j.tics.2019.04.013

Kanwisher, N., McDermott, J., & Chun, M. M. (1997). The fusiform face area: a module in human extrastriate cortex specialized for face perception. Journal of neuroscience, 17(11), 4302–4311. https://doi.org/10.1523/JNEUROSCI.17-11-04302.1997

Kaposvari, P., Kumar, S., & Vogels, R. (2018). Statistical learning signals in macaque inferior temporal cortex. Cerebral Cortex, 28(1), 250–266. https://doi.org/10.1093/cercor/bhw374

Kim, J. G., & Biederman, I. (2011). Where do objects become scenes?. Cerebral Cortex, 21(8), 1738–1746. https://doi.org/10.1093/cercor/bhq240

Kok, P., de Lange, F. (2015). Predictive Coding in Sensory Cortex. In: Forstmann, B., Wagenmakers, EJ. (eds) An Introduction to Model-Based Cognitive Neuroscience. Springer, New York, NY. https://doi.org/10.1007/978-1-4939-2236-9_11

Kourtzi, Z., & Kanwisher, N. (2001). Representation of perceived object shape by the human lateral occipital complex. Science, 293(5534), 1506–1509. https://doi.org/10.1126/science.1061133

Large, M. E., Aldcroft, A., & Vilis, T. (2007). Task-related laterality effects in the lateral occipital complex. Brain research, 1128, 130–138.

Lescroart, M. D., & Gallant, J. L. (2019). Human scene-selective areas represent 3D configurations of surfaces. Neuron, 101(1), 178–192. https://doi.org/10.1016/j.neuron.2018.11.004

Livne, T., & Bar, M. (2016). Cortical integration of contextual information across objects. Journal of cognitive neuroscience, 28(7), 948–958. https://doi.org/10.1162/jocn_a_00944

Malach, R., Reppas, J. B., Benson, R. R., Kwong, K. K., Jiang, H., Kennedy, W. A.,… & Tootell, R. B. (1995). Object-related activity revealed by functional magnetic resonance imaging in human occipital cortex. Proceedings of the National Academy of Sciences, 92(18), 8135–8139. https://doi.org/10.1073/pnas.92.18.8135

Malikovic, A., Amunts, K., Schleicher, A., Mohlberg, H., Kujovic, M., Palomero-Gallagher, N.,… & Zilles, K. (2016). Cytoarchitecture of the human lateral occipital cortex: mapping of two extrastriate areas hOc4la and hOc4lp. Brain Structure and Function, 221(4), 1877–1897. https://doi.org/10.1007/s00429-015-1009-8

McAndrews, M. P., Girard, T. A., Wilkins, L. K., & McCormick, C. (2016). Semantic congruence affects hippocampal response to repetition of visual associations. Neuropsychologia, 90, 235–242.

McLaren, D. G., Ries, M. L., Xu, G., & Johnson, S. C. (2012). A generalized form of context-dependent psychophysiological interactions (gPPI): a comparison to standard approaches. Neuroimage, 61(4), 1277–1286. https://doi.org/10.1016/j.neuroimage.2012.03.068

Oliva, A., & Torralba, A. (2007). The role of context in object recognition. Trends in Cognitive Sciences, 11(12), 520–527. https://doi.org/10.1016/j.tics.2007.09.009

Rao, R. P. N., & Ballard, D. H. (1999). Predictive coding in the visual cortex: A functional interpretation of some extra-classical receptive-field effects. Nature Neuroscience, 2(1), 79–87. https://doi.org/10.1038/4580

Rémy, F., Saint-Aubert, L., Bacon-Macé, N., Vayssière, N., Barbeau, E., & Fabre-Thorpe, M. (2013). Object recognition in congruent and incongruent natural scenes: A life-span study. Vision research, 91, 36–44. https://doi.org/10.1016/j.visres.2013.07.006

Rémy, F., Vayssière, N., Pins, D., Boucart, M., & Fabre-Thorpe, M. (2014). Incongruent object/context relationships in visual scenes: Where are they processed in the brain?. Brain and cognition, 84(1), 34–43. https://doi.org/10.1016/j.bandc.2013.10.008

Rémy, F., Vayssière, N., Saint-Aubert, L., Bacon-Macé, N., Pariente, J., Barbeau, E., & Fabre-Thorpe, M. (2020). Age effects on the neural processing of object-context associations in briefly flashed natural scenes. Neuropsychologia, 136, 107264.

Sawamura, H., Orban, G. A., & Vogels, R. (2006). Selectivity of neuronal adaptation does not match response selectivity: a single-cell study of the FMRI adaptation paradigm. Neuron, 49(2), 307–318. https://doi.org/10.1016/j.neuron.2005.11.028

Sobotka, S., & Ringo, J. L. (1994). Stimulus specific adaptation in excited but not in inhibited cells in inferotemporal cortex of macaque. Brain research, 646(1), 95–99. https://doi.org/10.1016/0006-8993(94)90061-2

Spaak, E., Peelen, M. V., & de Lange, F. P. (2022). Scene context impairs perception of semantically congruent objects. Psychological Science, 33(2), 299–313. https://doi.org/10.1177/09567976211032676

Summerfield, C., Egner, T., Greene, M., Koechlin, E., Mangels, J., & Hirsch, J. (2006). Predictive codes for forthcoming perception in the frontal cortex. Science, 314(5803), 1311–1314. https://doi.org/10.1126/science.1132028

Summerfield, C., Trittschuh, E. H., Monti, J. M., Mesulam, M., & Egner, T. (2008). Neural repetition suppression reflects fulfilled perceptual expectations. Nature neuroscience, 11(9), 1004–1006. https://doi.org/10.1038/nn.2163

Sutton, R. S., & Barto, A. G. (2018). Reinforcement learning: An introduction. MIT press.

Trapp, S., & Bar, M. (2015). Prediction, context, and competition in visual recognition. Annals of the New York Academy of Sciences, 1339(1), 190–198.

van Kesteren, M. T., Rignanese, P., Gianferrara, P. G., Krabbendam, L., & Meeter, M. (2020). Congruency and reactivation aid memory integration through reinstatement of prior knowledge. Scientific Reports, 10(1), 1–13.

Weilnhammer, V., Stuke, H., Hesselmann, G., Sterzer, P., & Schmack, K. (2017). A predictive coding account of bistable perception-a model-based fMRI study. PLoS computational biology, 13(5), e1005536. https://doi.org/10.1371/journal.pcbi.1005536

Weiner, K. S., & Zilles, K. (2016). The anatomical and functional specialization of the fusiform gyrus. Neuropsychologia, 83, 48–62. https://doi.org/10.1016/j.neuropsychologia.2015.06.033

Weiner, K. S., Natu, V. S., & Grill-Spector, K. (2018). On object selectivity and the anatomy of the human fusiform gyrus. NeuroImage, 173, 604–609. https://doi.org/10.1016/j.neuroimage.2018.02.040

Whitfield-Gabrieli, S., & Nieto-Castanon, A. (2012). Conn: a functional connectivity toolbox for correlated and anticorrelated brain networks. Brain connectivity, 2(3), 125–141. https://doi.org/10.1089/brain.2012.0073

